# BRM preserves sinusoidal niche resilience through Jag2–Notch signaling during hematopoietic regeneration and aging

**DOI:** 10.64898/2026.07.03.736310

**Authors:** Haruka Suzuki, Hiroki Miyachi, Natsuka Yamada, Hiroki Nishikawa, Shuhei Kuno, Hiromi Endoh, Tatsuro Shiina, Ryo Yoshikawa, Shun Uemura, Atsushi Iwama, Toshio Suda, Ryo Nitta, Eriko Nitta

## Abstract

During aging and stress, remodeling of the bone marrow (BM) microenvironment compromises hematopoietic stem cell (HSC) maintenance, contributing to extrinsic HSC aging. Although niche-derived Notch signaling is essential for hematopoietic regeneration following myelosuppressive injury, the upstream chromatin regulatory mechanisms governing this stress-responsive signaling remain poorly understood. Here, we identify the chromatin remodeler BRM as a critical regulator of the BM sinusoidal niche integrity and hematopoietic regeneration. Using reciprocal BM transplantation, we demonstrate that a Brm-deficient microenvironment impairs hematopoietic reconstitution and imposes an aging-like myeloid bias characterized by expansion of granulocyte-monocyte progenitors. Following 5-fluorouracil (5-FU)-induced myelosuppression, BrmKO mice exhibit defective sinusoidal regeneration accompanied by endothelial degeneration. Mechanistically, BRM deficiency compromises Notch signaling at two levels: it reduces the Jag2 ligand pool through persistent depletion of sinusoidal endothelial cells (SECs) and impaired regenerative expansion of Jag2-producing LepR^+^ stromal cells, while also impairing stress-induced Notch2 expression and downstream Hes1 activation in SECs. These defects are accompanied by impaired sinusoidal regeneration and failure to restore normal HSC positioning relative to the sinusoidal vasculature. Notably, the Brm-deficient microenvironment recapitulates key features of physiological niche aging, while BRM expression declines with age in SECs. Collectively, our findings identify BRM as a key regulator of BM niche regenerative resilience and suggest that age-associated BRM decline contributes to niche dysfunction and hematopoietic aging.

**Key Points:**

- Microenvironmental BRM sustains hematopoietic reconstitution and prevents aging-like myeloid bias.
- BRM preserves Jag2 availability and Notch2 signaling during stress, supporting sinusoidal regeneration and normal HSC positioning.

## Introduction

Hematopoietic stem cells (HSCs) sustain the lifelong production of blood and immune cells through their capacity for self-renew and multilineage differentiation^1–6^. This process is critically governed by the bone marrow (BM) microenvironment, or niche, which provides structural and molecular cues essential for HSC maintenance^7–14^. The niche comprises diverse non-hematopoietic components, including arteriolar endothelial cells (AECs), sinusoidal endothelial cells (SECs), mesenchymal stromal cells (MSCs), and osteoblast-lineage cells^15,16^. Among these components, SECs and perivascular stromal cells constitute the sinusoidal niche^17–23^, a specialized microenvironment that regulates HSC quiescence, localization, and regenerative capacity and is preferentially preserved during aging to sustain residual HSC function^24^. With aging and chronic stress, remodeling of the BM microenvironment—characterized by vascular alterations and stromal dysfunction—profoundly impairs HSC homeostasis and contributes functional hematopoietic decline^5,25–32^. However, the upstream mechanisms that preserve niche integrity and regenerative capacity during stress and aging remain poorly understood^33,34^.

A critical communication pathway between the niche and HSCs is the Notch signaling, which endothelial- and mesenchymal-derived ligands regulate adult BM homeostasis and regeneration^32,35,36^. Specifically, the Notch ligand Jagged-2 (Jag2), predominantly expressed by SECs and leptin receptor (LepR)-expressing stromal cells, contributes to the spatial organization of HSCs within the supportive vascular niche^24,37,38^. Beyond direct niche-to-HSC crosstalk, recent evidence indicates that autocrine/paracrine endothelial Notch signaling is required for vascular regeneration following myelosuppressive injury, thereby supporting hematopoietic recovery^39^. Thus, dynamic regulation of Jag2–Notch signaling within the BM microenvironment appears critical for post-injury niche repair^24,37^, yet the upstream mechanisms that coordinate this stress-responsive signaling remain unknown.

Recent studies, including ours, have identified the chromatin remodeler BRM (SMARCA2) as a key regulator of stress hematopoiesis. BRM preserves HSC function during regenerative stress and limits aging-associated hematopoietic decline by maintaining stress-responsive chromatin accessibility^40^. These findings raised the possibility that BRM may also coordinate hematopoietic stress responses through regulation of the BM microenvironment. We therefore hypothesized that BRM contributes to the regenerative capacity of the sinusoidal niche during hematopoietic stress.

Here, we identify BRM as a key regulator of sinusoidal niche resilience during hematopoietic regeneration and aging. Using Brm-deficient mice and bone marrow transplantation, we show that microenvironmental BRM supports hematopoietic reconstitution and preserves sinusoidal vascular integrity following myelosuppressive injury. Mechanistically, BRM maintains the Jag2 ligand pool by preserving Jag2-producing niche populations while supporting stress-induced Notch2 expression and downstream Hes1 activation in SECs. BRM deficiency is accompanied by defective sinusoidal regeneration, altered HSC positioning, and aging-like myeloid bias, and BRM expression declines in SECs during physiological aging. Together, these findings identify BRM-dependent chromatin remodeling as a mechanism that supports regenerative resilience of the BM niche and links impaired stress adaptation to hematopoietic aging.

## Methods

### Mice and 5-FU Treatment Model

Brm-deficient (BrmKO) mice were provided by Dr. Tadao Arinami and Dr. Yoshimi Iijima (University of Tsukuba) and maintained under specific pathogen-free (SPF) conditions. Ly5.1 (CD45.1) congenic mice were purchased from Sankyo Labo Service. Male and female mice aged 8-12 weeks were used as young adult controls, whereas mice aged >80 weeks were used as aged animals. For myelosuppressive injury, mice received a single intraperitoneal injection of 5-fluorouracil (5-FU; Kyowa Kirin) at 250 mg/kg and were analyzed at days 4, 30, and 120 after treatment, representing acute injury and subsequent recovery phases. Untreated mice were analyzed as steady-state controls. All animal protocols were approved by the Kobe University Institutional Animal Care and Use Committee (Approval No. P-221104).

### Isolation of Bone Marrow Niche Cells

To isolate bone marrow niche cells, bone marrow was flushed from the right femurs and tibia with Accumax (Innovative Cell Technologies), followed by rotation at 25 rpm and 25℃ for 45-60 minutes. Cells were resuspended in phosphate-buffered saline (PBS, pH 7.4) containing 0.5% bovine serum albumin (BSA), and red blood cells were lysed in ammonium chloride-based lysis buffer for 3 minutes. Cells were incubated with Fc block and biotinylated anti-Leptin Receptor (LepR) antibody (R&D Systems) at 4℃ for 30 minutes, followed by staining with streptavidin and antibodies against Ter119, Jagged2, CD31, Sca-1, CD45.2, CD140a, and CD105. SECs were defined as CD45^−^ Ter119^−^CD31^+^Sca-1^+^CD105^+^, LepR⁺ stromal cells as CD45^−^Ter119^−^CD31^−^CD140a ^+^LepR^+^, and arteriolar endothelial cells (AECs) as CD45^−^Ter119^−^CD31^+^Sca-1^+^CD105^−^, with propidium iodide (PI) used to exclude dead cells. Cells were sorted using a FACSMelody (BD Biosciences) or analyzed using a BD LSRFortessa, and data were processed with FlowJo v10.10.1. Details of antibodies used are summarized in Supplemental Table 1.

### Bone Marrow Transplantation into WT and BrmKO Recipients

WT and BrmKO recipient mice (CD45.2, 8-12 weeks old) were lethally irradiated with a total dose of 8.5 Gy administered in two fractions (4.5 and 4.0 Gy, 3 hours apart) using an X-ray irradiator (MBR-1605R-BE, Hitachi; 150 kV, 4 mA). 2.0×10⁵ whole bone marrow cells from young WT donors (CD45.1) were injected via the tail vein using a 28G needle. Donor chimerism (CD45.1^+^) in peripheral blood (PB) was monitored every 4 weeks, and recipient BM was analyzed 16 weeks after transplantation.

### PB Chimerism Analysis

At 4, 8, 12, and 16 weeks after transplantation, PB was collected from the tail vein into EDTA-coated capillary tubes. Following red blood cell lysis, leukocytes were stained with antibodies against CD45.1 and CD45.2 and analyzed on a BD LSRFortessa.

### Analysis of Hematopoietic Stem and Progenitor Cells

Bone marrow was flushed from femurs and tibiae with PBS, followed by red blood cell lysis. Cells were stained with the indicated antibodies and analyzed on a BD LSRFortessa. Donor-derived cells were identified as CD45.1⁺ cells. Within Lineage^−^ Sca-1^+^c-Kit^+^ (LSK) cells, HSCs were defined as CD34^−^Flt3^−^CD48^−^CD150^+^; MPP1 as CD34^+^Flt3^−^CD48^−^CD150^+^; MPP2 as CD34^+^Flt3^−^CD48^+^CD150^+^; MPP3 as CD34^+^Flt3^−^ CD48^+^CD150^−^; and MPP4 as CD34^+^Flt3^+^CD48^−^CD150^−^. Within Lineage^−^Sca-1^−^c-Kit^+^ cells, GMPs were defined as CD34^+^FcγR^+^, MEPs as CD34^−^FcγR^−^, and CMPs as CD34^+^FcγR^−^.

### Histological Analysis (H&E Staining)

Femurs were fixed in 4% paraformaldehyde (PFA) in PBS at 4℃ for 2–3 days and decalcified in EDTA-4Na at 4℃ for 7 days. Samples were then dehydrated, embedded in paraffin, sectioned, and stained with hematoxylin and eosin (H&E). Images were acquired using a BZ-X700 microscope (Keyence) with 4×, 20×, and 40× objectives.

### Sternal Whole-Mount Immunostaining

Sterna were cut longitudinally to expose the bone marrow cavity. Tissues were fixed in 4% PFA in phosphate buffer at room temperature for 15-30 minutes and washed with Tris-buffered saline containing 1 mM CaCl_2_. Samples were blocked and permeabilized in 10% normal donkey serum containing 0.5% Triton X-100 for 1 hour at room temperature, followed by incubation at 4℃ for 24 hours with anti-CD150 (1:25), anti-CD144 (1:50), rabbit anti-Jagged2 (1:100), and lineage cocktail antibodies, and then incubated with the appropriate fluorophore-conjugated secondary antibodies for 2 hours at room temperature.

### 3D Confocal Imaging and Spatial Distance Quantification

Three-dimensional images were acquired using an FLUOVIEW FV4000 confocal microscope (EVIDENT/Olympus) with 10× or 30× objectives and processed with ZEN desk (Carl Zeiss) and Fiji/ImageJ (v2.0.2). The shortest three-dimensional distance (μm) between each HSC (Lineage^−^CD150^+^) and the surface of the nearest CD144^+^ sinusoidal vessel was measured^41^.

### Quantitative Analysis of CD144 (VE-Cadherin) and Jagged2 Fluorescence Intensity

Z-stack images were acquired using identical settings across groups, and maximum-intensity projections were generated in Fiji/ImageJ. CD144+ (VE-Cadherin) sinusoidal vessel regions of interest were defined by thresholding, and mean fluorescence intensity (mean gray value) was measured. For Jag2, regions of interest (ROIs) of identical size were defined in each image, and the raw integrated density (RawIntDen) of each ROI was measured. Background signal was estimated from ten background ROIs per image as the mean RawIntDen per unit area, multiplied by the Jag2 ROI area, and subtracted from the RawIntDen of each Jag2 ROI. The background-corrected value was then divided by the ROI area and is presented as relative fluorescence intensity (arbitrary units).

### Transmission Electron Microscopy (TEM) Analysis

Bone marrow was gently flushed from femurs and tibiae into ice-cold 0.1 M phosphate buffer and fixed in 2.5% glutaraldehyde in 0.1 M phosphate buffer for 2 hours at room temperature. Samples were postfixed in 1% osmium tetroxide for 2 hours at 4℃, dehydrated through a graded ethanol series, embedded in Epon 812 via propylene oxide, and polymerized at 60℃ for 48 hours. Ultrathin sections (70 nm) were cut with a diamond knife, collected on grids, stained with uranyl acetate for 15 minutes and lead solution for 3 minutes, and examined using a JEM-100 electron microscope (JEOL) for ultrastructural analysis.

### Quantitative Real-Time PCR (qPCR) Analysis

SECs were sorted directly into the lysis buffer of an RNeasy Plus Micro Kit (Qiagen), and total RNA was extracted with genomic DNA elimination. cDNA was synthesized using SuperScript III (Invitrogen). Quantitative PCR was performed on a QuantStudio 5 or LightCycler 480 System using THUNDERBIRD Next SYBR qPCR Mix (TOYOBO), and gene expression was normalized to *Gapdh*. *Smarca2 (Brm*) was measured using a TaqMan Gene Expression Assay (Mm01325629_m1) and TaqMan Fast Advanced Master Mix and normalized to *Actb* (Mm00607939_s1). Relative expression was calculated using the 2⁻ΔΔCt method. Primer sequences are listed below:

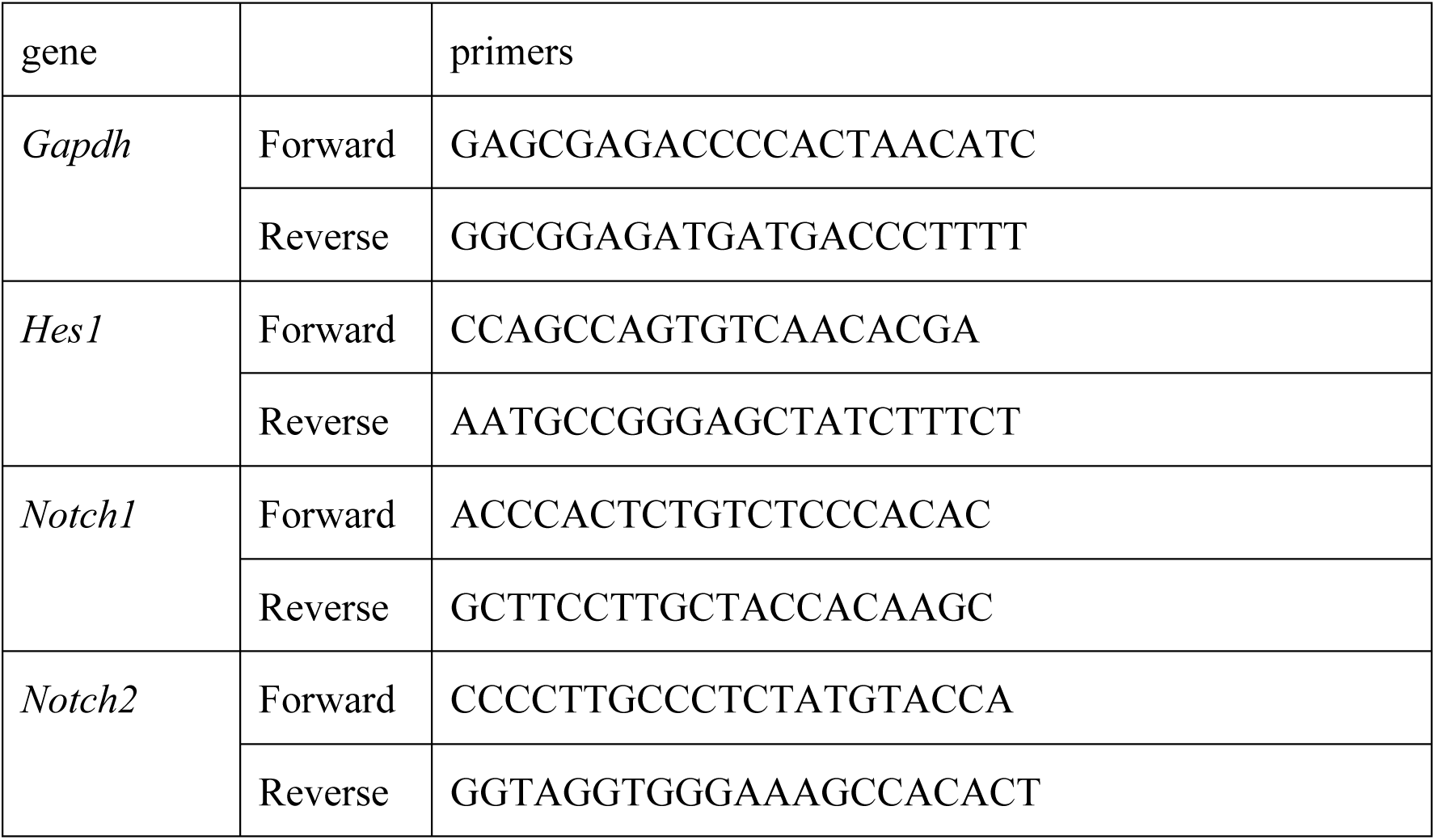

### Analysis of public RNA-seq data

BRM expression in bone marrow niche populations of 2-month-old adult and 2-year-old aged mice was analyzed using the publicly available RNA-seq dataset GSE133922^42^, using the normalized count matrix deposited by the original authors. Expression was compared between age groups within each cell population using a two-tailed Welch’s *t* test without correction for multiple comparisons.

### Statistical Analysis and Data Visualization

Statistical analyses and data visualization were performed using an in-house Python (v3.11.5) pipeline with Pandas, NumPy, SciPy, and Matplotlib. Data are presented as mean ± SD or SEM, as indicated in the figure legends. Comparisons between two groups were performed using a two-tailed Mann–Whitney U test, except for the reanalysis of public RNA-seq data, in which a two-tailed Welch’s t test was used. Statistical significance was defined as *P* < 0.05 (\**P* < 0.05, \*\**P* < 0.01, \*\*\**P* < 0.001, \*\*\*\**P* < 0.0001; ns, *P* ≥ 0.05). n indicates independent mice or experiments, as specified in the figure legends.

## Results

### Microenvironmental BRM preserves hematopoietic reconstitution and prevents myeloid bias

To determine whether BRM is required for the hematopoietic supportive function of the bone marrow (BM) microenvironment, we transplanted wild-type (WT; Ly5.1) whole BM cells into lethally irradiated WT (Ly5.2) or Brm-deficient (BrmKO; Ly5.2) recipient mice, thereby selectively assessing the contribution of recipient-derived BRM to hematopoietic reconstitution (Figure 1A).

**Figure 1.**
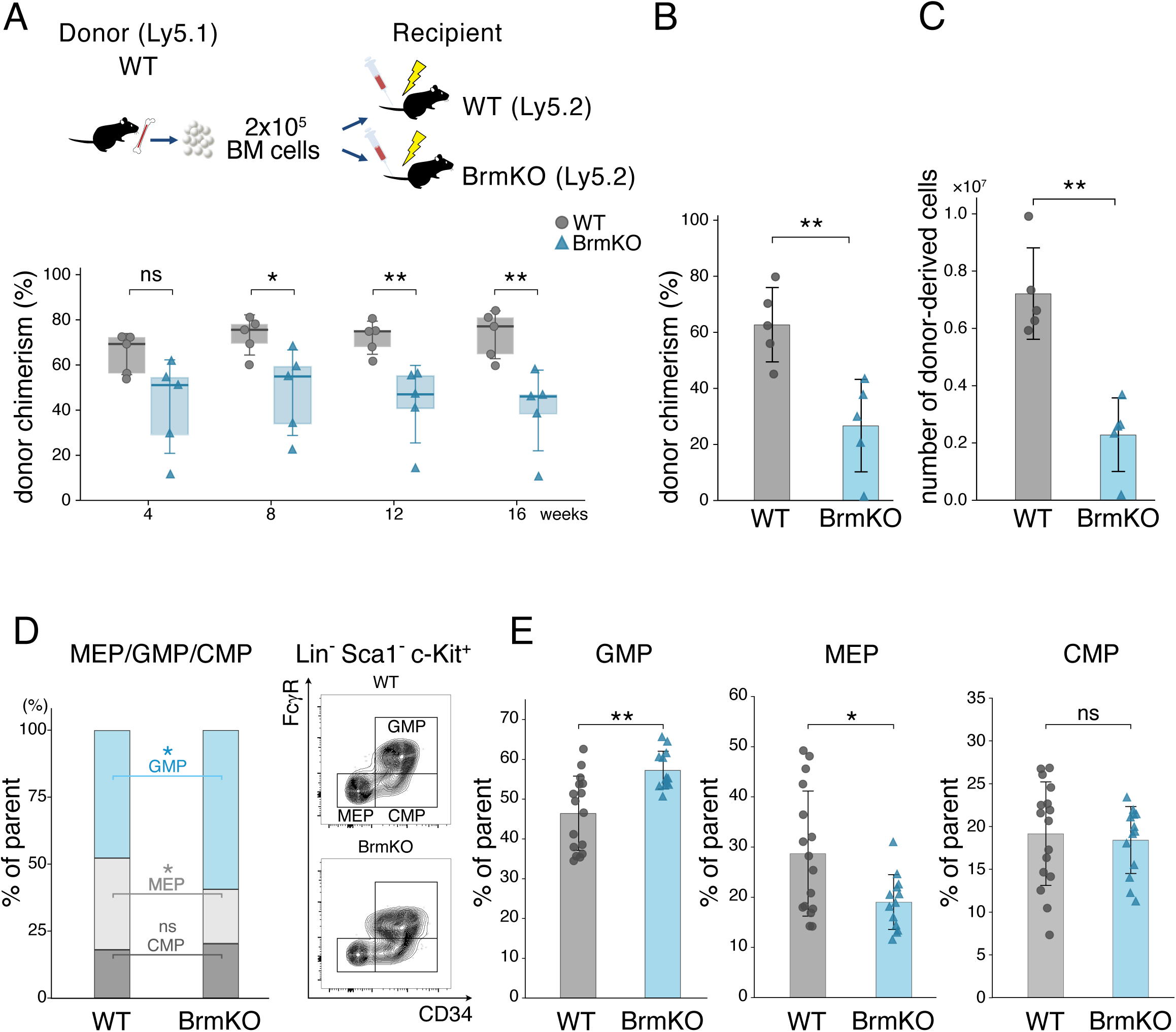
Microenvironmental BRM preserves HSC reconstitution and prevents myeloid bias. (A) Schematic representation of the bone marrow transplantation (BMT) model (upper) and donor-derived chimerism (CD45.1⁺) in the peripheral blood (PB) of WT and BrmKO recipient mice (CD45.2⁺) (lower) (WT, *n* = 5; BrmKO, *n* = 5; representative of 3 independent experiments). (B) Donor-derived chimerism (CD45.1⁺) in the bone marrow (BM) at 16 weeks after transplantation (WT, *n* = 5; BrmKO, *n* = 5). (C) Absolute number of donor-derived (CD45.1⁺) live BM cells (WT, *n* = 5; BrmKO, *n* = 5), calculated by multiplying the total number of harvested BM cells by the frequency of CD45.1⁺ cells. (D) Quantification (left) and representative flow cytometry plots (right) of donor-derived myeloid progenitor populations in the BM of recipient mice. The relative proportions of GMPs, MEPs, and CMPs within the donor-derived lineage⁻Sca-1⁻c-Kit⁺ compartment were normalized to 100% (WT, n = 11; BrmKO, n = 10). (E) Frequencies of donor-derived GMPs, MEPs, and CMPs in the BM of recipient mice (WT, *n* = 16; BrmKO, *n* = 13). Data are pooled from at least 3 independent experiments and presented as mean ± SD. *P* values were calculated using a two-tailed Mann-Whitney U test. ns, not significant; \**P* < 0.05; \*\**P* < 0.01.

Donor-derived (CD45.1⁺) chimerism in peripheral blood (PB) progressively declined in BrmKO recipients from 4 to 16 weeks after transplantation (Figure 1A). Consistent with this finding, donor chimerism in the BM remained significantly lower at 16 weeks (Figure 1B), accompanied by a reduced absolute number of donor-derived BM cells (Figure 1C), indicating impaired donor-derived hematopoietic reconstitution. Despite this reduction, the frequencies of donor-derived phenotypic HSCs and multipotent progenitors (MPP1–4) were comparable between genotypes (Supplementary Figure 1A, B). In contrast, myeloid progenitor populations exhibited a marked imbalance in BrmKO recipients, characterized by an increased proportion of granulocyte–monocyte progenitors (GMPs) and a reduced proportion of megakaryocyte–erythroid progenitors (MEPs), whereas common myeloid progenitors (CMPs) remained unchanged (Figure 1D, E).

Together, these findings demonstrate that microenvironmental BRM is required to sustain hematopoietic reconstitution and maintain balanced myeloid progenitor composition following transplantation.

### BRM differentially regulates vascular niche populations during regeneration and aging

To assess the contribution of BRM to bone marrow niche regeneration independently of irradiation and its relevance to physiological aging, we employed an irradiation-free 5-FU model^43^. Analysis of a publicly available RNA-seq datasets showed that *Brm* expression declines with age in SECs (Supplementary Figure 1C) ^42^, suggesting a potential link between age-associated BRM decline and niche dysfunction. We therefore quantified the three major vascular and perivascular niche populations—SECs, AECs, and LepR⁺ stromal cells—during 5-FU-induced regeneration and physiological aging by flow cytometry.

SEC frequencies were significantly reduced in BrmKO mice in steady state and remained lower throughout hematopoietic regeneration (days 4 and 30 after 5-FU treatment; Figure 2A, D). Consistent with previous reports, SEC frequencies declined during physiological aging in WT mice; however, they remained significantly lower in aged BrmKO mice than in aged WT mice (Figure 2A, D).

**Figure 2.**
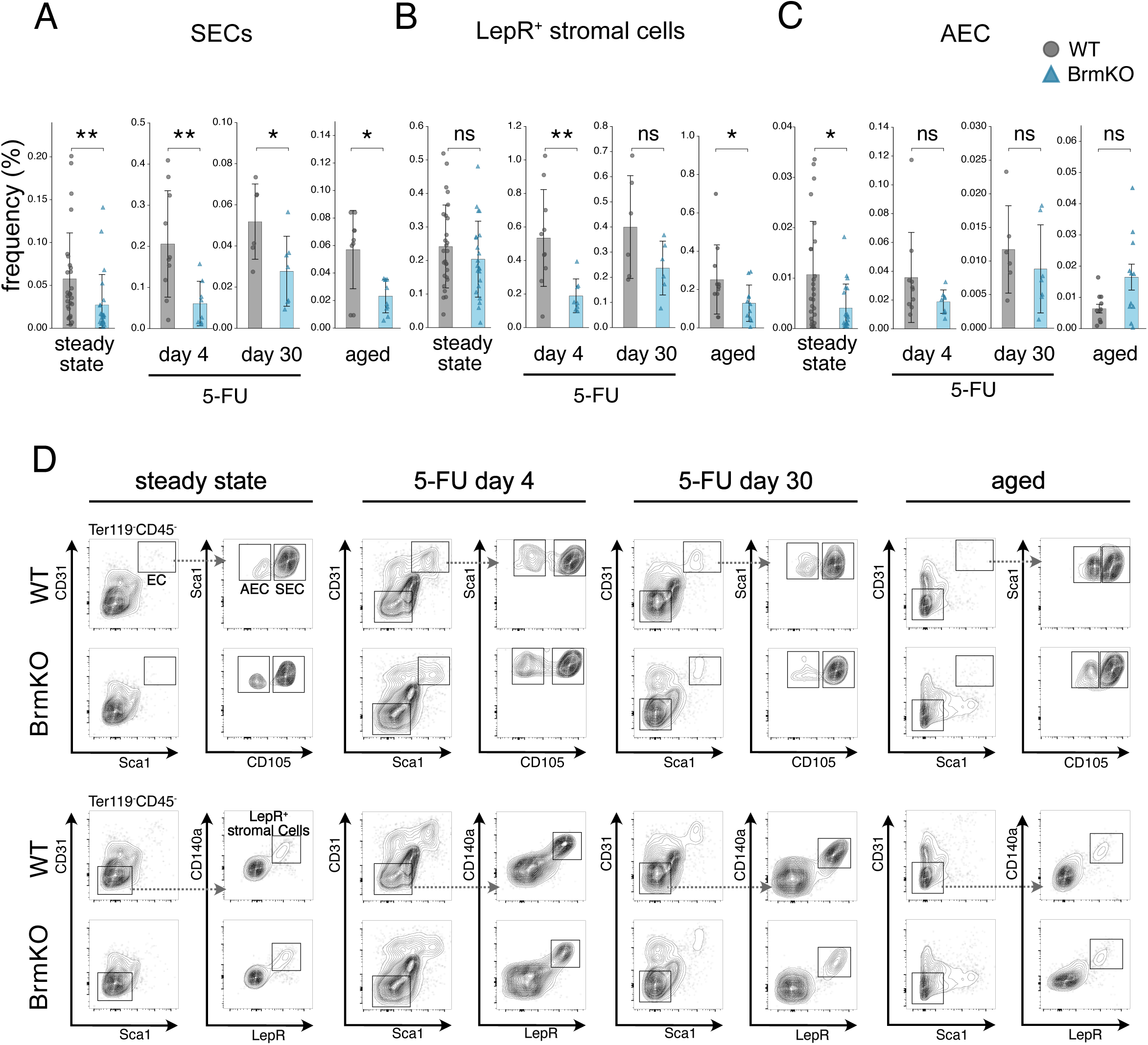
BRM differentially regulates vascular niche populations during regeneration and aging. (A-C) Flow cytometric quantification of the frequencies of SECs (A), LepR⁺ stromal cells (B), and AECs (C) at steady state, days 4 and 30 after 5-FU treatment, and in aged mice (*n* = 6–28 mice per group across conditions). Frequencies are expressed as percentages of the Ter119⁻ fraction. (D) Representative flow cytometry gating plots for SECs, AECs, and LepR⁺ stromal cells. Data are pooled from at least 3 independent experiments and presented as mean ± SD. *P* values were calculated using a two-tailed Mann-Whitney U test. ns, not significant; \**P* < 0.05; \*\**P* < 0.01.

In contrast, LepR⁺ stromal cells exhibited a transient regenerative defect. Their frequencies were comparable between genotypes at steady state but were significantly reduced in BrmKO mice during the early regenerative phase (day 4), with no significant difference between genotypes by day 30. LepR⁺ stromal cell frequencies were also significantly reduced in aged BrmKO mice compared with aged WT mice (Figure 2B, D). AECs were modestly reduced in BrmKO mice at steady state but showed no genotype-dependent differences during regeneration or physiological aging (Figure 2C, D).

Together, these findings demonstrate that BRM differentially regulates bone marrow niche populations, primarily by maintaining the SEC compartment during homeostasis and regeneration, while supporting the regenerative response of LepR⁺ stromal cells.

### BRM preserves sinusoidal vascular integrity during regeneration and aging

We next examined bone marrow regeneration histologically by H&E staining and found that BrmKO mice exhibited impaired sinusoidal regeneration accompanied by marked sinusoidal dilation, particularly at day 4 after 5-FU administration (Supplementary Figure 2A).

Whole-mount imaging of CD144⁺ sinusoids further revealed enlarged sinusoidal vessels in BrmKO mice even at steady state (Figure 3A, B). Following 5-FU treatment, WT mice progressively restored a fine reticular sinusoidal network by day 30, whereas BrmKO mice exhibited persistent sinusoidal ectasia that remained evident through day 120. Similar sinusoidal abnormalities were observed in BrmKO recipients transplanted with wild-type hematopoietic cells (Figure 3A, B), further supporting a microenvironmental origin of the phenotype. Comparable abnormalities were also observed in physiologically aged BrmKO mice (Figure 3A, B).

**Figure 3.**
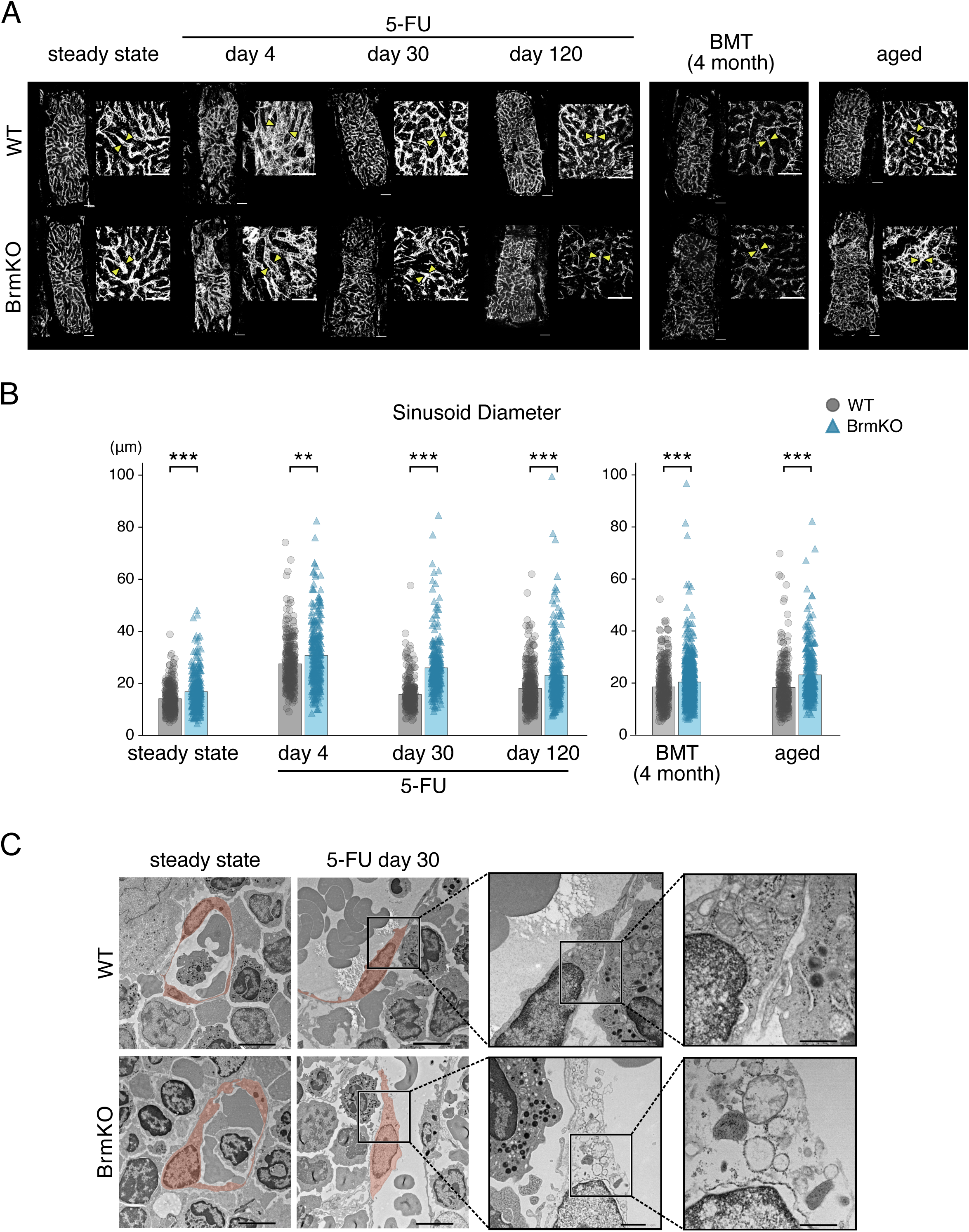
BRM preserves sinusoidal vascular integrity during regeneration and aging. (A) Representative whole-mount immunofluorescence images of CD144⁺ sinusoids in the sternal bone marrow of WT and BrmKO mice at steady state, after 5-FU treatment (days 4, 30, and 120), 4 months after BMT, and in aged mice. Magnified views are shown to the right of each panoramic image. Paired yellow arrowheads indicate representative sinusoidal vessels. Scale bars, 200 μm. (B) Quantification of sinusoidal vessel diameters under the conditions shown in (A). At least 100 vessel diameters per mouse were measured from sternal whole-mount immunofluorescence images (*n* = 3 mice per group for each condition). (C) Representative transmission electron microscopy (TEM) images of SECs at steady state and day 30 after 5-FU treatment. SECs lining the sinusoids are pseudocolored in orange. Scale bars, 5 μm, 1 μm, and 500 nm. Data are pooled from at least 3 independent experiments and presented as mean ± SD. *P* values were calculated using a two-tailed Mann-Whitney U test. ns, not significant; \*\**P* < 0.01; \*\*\**P* < 0.001.

To define the cellular basis of these structural abnormalities, we examined the ultrastructure of bone marrow niche cells by transmission electron microscopy (TEM). At steady state, SEC morphology was largely comparable between genotypes. In contrast, at day 30 after 5-FU treatment, BrmKO SECs exhibited dilation of the nuclear envelope, organelle disruption, and cytoplasmic rarefaction (Figure 3C). These ultrastructural abnormalities were not observed in perivascular stromal cells (Supplementary Figure 2B). Moreover, endothelial cell–cell junctions and CD144 (VE-cadherin) expression were preserved in BrmKO mice at steady state and day 30 after 5-FU treatment (Supplementary Figure 2C, D).

Together, these findings indicate that defective regeneration of the BrmKO sinusoidal niche is associated with impaired cellular integrity of SECs rather than overt disruption of endothelial junctions.

### BRM sustains the Jag2 ligand pool by maintaining Jag2-producing niche populations

Given the emerging role of endothelial Jag2 in regulating hematopoietic regeneration and age-associated HSC maintenance^37,38^, we next examined Jag2 expression in the bone marrow by whole-mount immunofluorescence (Figure 4A). Jag2 levels were modestly reduced in BrmKO mice at steady state. More strikingly, Brm deficiency markedly impaired the increase in Jag2 during regeneration following 5-FU treatment (days 4–30) and bone marrow transplantation. Furthermore, the physiological increase in Jag2 observed during aging was significantly attenuated in BrmKO mice (Figure 4A).

**Figure 4.**
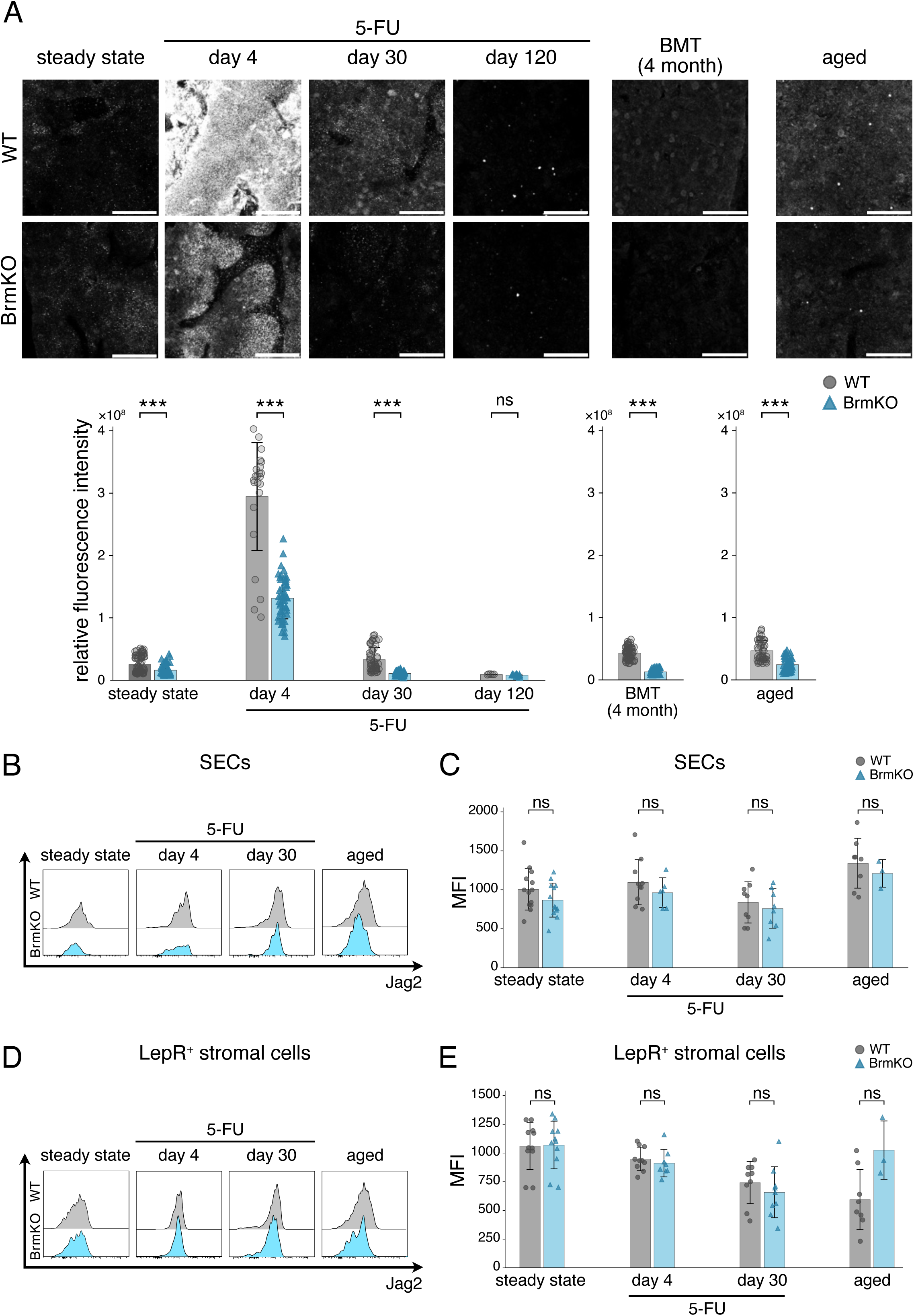
BRM sustains Jag2 availability by preserving Jag2-producing niche populations. (A) Representative whole-mount immunofluorescence images (upper) and quantification of relative Jag2 fluorescence intensity (lower) in the sternal bone marrow at steady state, after 5-FU treatment (days 4, 30, and 120), after BMT, and in aged mice. Scale bar, 200 μm. n = 3 mice per group; each dot represents one ROI. Data are presented as mean ± SD. (B) Representative flow cytometric histogram overlays of Jag2 expression in SECs. (C) Quantification of Jag2 median fluorescence intensity (MFI) in SECs (n = 3–13 per group). (D) Representative flow cytometric histogram overlays of Jag2 expression in LepR⁺ stromal cells. (E) Quantification of Jag2 MFI in LepR⁺ stromal cells (n = 3–11 per group). Data are pooled from at least 3 independent experiments and shown as mean ± SD. *P* values were calculated using a two-tailed Mann-Whitney U test. ns, not significant; \*\*\**P* < 0.001.

To determine the cellular basis of this reduction, we examined Jag2 expression in major bone marrow niche populations. In SECs, histogram analysis revealed a subtle leftward shift in Jag2 fluorescence in BrmKO mice, although median fluorescence intensity did not differ significantly between genotypes (Figure 4B, C). Jag2 expression in LepR⁺ stromal cells was also comparable between genotypes under both homeostatic and regenerative conditions (Figure 4D, E). In contrast, BrmKO mice exhibited a persistent reduction in SEC frequency and an impaired regenerative response of LepR⁺ stromal cells (Figure 2A, B). Collectively, these findings suggest that the reduction in Jag2 within BrmKO bone marrow primarily reflects quantitative depletion of Jag2-producing niche cells—through persistent loss of SECs and impaired regenerative expansion of LepR⁺ stromal cells—rather than by substantial downregulation of Jag2 expression within individual niche cells.

AECs expressed uniformly low levels of Jag2 under all conditions, with no significant difference between genotypes (Supplementary Figure 3A). CD45⁺ hematopoietic cells transiently upregulated Jag2 during early regeneration; however, neither the frequency of Jag2⁺CD45⁺ cells nor Jag2 expression in CD45⁺ cells differed between WT and BrmKO mice, arguing against a major contribution of the hematopoietic compartment to the BRM-dependent reduction in bone marrow Jag2 (Supplementary Figure 3B-D).

### BRM sustains Notch signaling in SECs during regeneration and aging

We next examined the expression of the Jag2 receptors *Notch1* and *Notch2* in FACS-isolated SECs. At steady state, expression of both receptors was comparable between WT and BrmKO mice. During regeneration following 5-FU treatment, however, stress-induced *Notch2* upregulation was significantly impaired in BrmKO SECs, whereas *Notch1* expression was not similarly affected (Figure 5A, B). Likewise, the age-associated increases in both *Notch1* and *Notch2* expression were attenuated in SECs from aged BrmKO mice. Notably, *Notch1* expression increased during the later recovery phase after 5-FU treatment despite persistent impairment of *Notch2* expression (Figure 5B).

**Figure 5.**
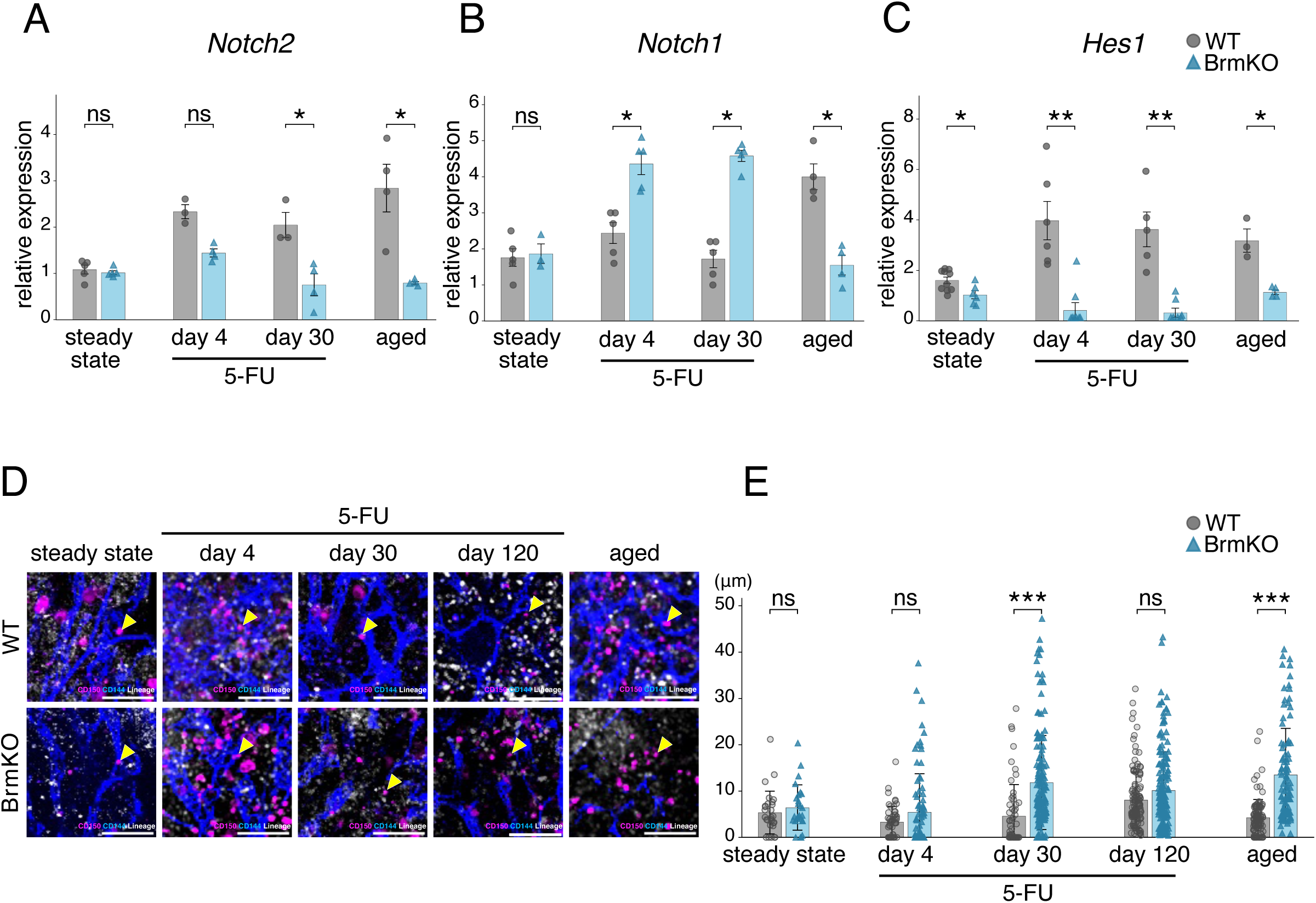
BRM sustains Notch signaling to preserve HSC positioning during sinusoidal regeneration and aging. (A-C) Reverse transcription quantitative PCR analysis of *Notch2* (A), *Notch1* (B), and *Hes1* (C) mRNA expression in primary SECs isolated via flow cytometry at steady state, after 5-FU treatment (days 4 and 30), and in aged mice (*n* = 3–5 per group). Expression levels were normalized to *Gapdh*, and relative fold changes were calculated using the 2⁻ΔΔCt method relative to steady-state WT SECs. (D) Representative whole-mount immunofluorescence images of sternal bone marrow showing CD150⁺Lineage⁻ HSCs (magenta; arrowheads) and CD144⁺ sinusoids (blue) in WT and BrmKO mice at steady state, at days 4, 30, and 120 after 5-FU treatment, and in aged mice. Scale bar, 100 μm. (E) Quantification of the shortest three-dimensional distance from individual HSCs to the nearest sinusoidal vessel wall under the indicated conditions (*n* = 3 mice per group). Each dot represents an individual HSC. Data are pooled from at least 3 independent experiments and presented as mean ± SEM (A–C) and mean ± SD (E). *P* values were calculated using a two-tailed Mann-Whitney U test. ns, not significant; \**P* < 0.05 and \*\*\**P* < 0.001.

Consistent with the reduced Jag2 availability and impaired *Notch2* induction, expression of the downstream Notch target *Hes1* was significantly reduced in BrmKO SECs. Although *Hes1* expression was only modestly reduced at steady state, its induction during regeneration was markedly impaired, particularly at day 4 after 5-FU treatment, and remained reduced through day 30 (Figure 5C). *Hes1* expression was also consistently reduced in SECs from aged BrmKO mice, recapitulating the signaling defect observed during regeneration.

Together, these findings indicate that BRM is required for robust Notch signaling in SECs during regenerative stress and aging, characterized by stress-responsive *Notch2* induction and downstream *Hes1* activation.

### BRM preserves HSC positioning during sinusoidal regeneration and physiological aging

To determine whether defective sinusoidal regeneration alters the spatial organization of the HSC niche, we measured the distance between CD150⁺Lineage⁻ HSCs and CD144⁺ sinusoidal vessels in whole-mount sternal bone marrow preparations (Figure 5D, E). At steady state, HSC-to-sinusoid distances were comparable between WT and BrmKO mice. During the early regenerative phase (day 4 after 5-FU treatment), BrmKO mice exhibited a trend toward increased HSC-to-sinusoid distance, although the difference did not reach statistical significance. By day 30, however, HSCs in BrmKO mice were located significantly farther from the sinusoidal vasculature than those in WT controls. At day 120, the difference was no longer statistically significant, although the trend toward greater HSC-to-sinusoid distance remained in BrmKO mice. A similar increase in HSC-to-sinusoid distance was also observed in aged BrmKO mice (Figure 5D, E).

Together, these findings indicate that BRM supports the restoration of normal HSC positioning during sinusoidal niche regeneration and maintenance of HSC spatial organization during aging.

## Discussion

In this study, we identify BRM as a critical regulator of the regenerative resilience of the bone marrow niche. Rather than simply maintaining the niche at steady state, BRM becomes particularly important when the microenvironment is challenged by myelosuppressive stress, supporting restoration of the sinusoidal vasculature and subsequent hematopoietic recovery. Consistent with this concept, a BRM-deficient microenvironment was sufficient to impair reconstitution of WT hematopoietic cells and impose a myeloid-biased progenitor composition, indicating that loss of BRM-dependent niche function can promote features of hematopoietic aging independently of HSC-intrinsic BRM deficiency. Thus, BRM-dependent chromatin remodeling may provide the niche with the capacity to adapt to injury and restore hematopoietic homeostasis.

A key mechanistic insight from our study is that BRM supports Jag2–Notch signaling at both the signal-producing and signal-receiving sides of the sinusoidal niche. The reduction in the bone marrow Jag2 pool in BrmKO mice was associated primarily with persistent loss of SECs and impaired regenerative response of Jag2-producing LepR⁺ stromal cells, rather than substantial suppression of Jag2 expression within individual niche cells. In parallel, BRM-deficient SECs failed to efficiently induce *Notch2* and its downstream target *Hes1* during regeneration. *Notch1* expression was relatively preserved and increased during the later recovery phase, potentially reflecting a compensatory response to impaired *Notch2* signaling; however, this increase was not accompanied by restoration of *Hes1* expression. Rather than constituting a simple linear BRM–Jag2 pathway, therefore, BRM appears to coordinate both the availability of an extracellular regenerative signal and the transcriptional competence of endothelial cells to respond to regenerative cues.

This model is consistent with established functions of endothelial Notch signaling in vascular homeostasis and regeneration. Under steady-state conditions, endothelial Hes1 has been implicated in regulating HSC quiescence through control of p27 expression^44^, whereas acute Notch–Hes1 activation following myelosuppressive injury protects endothelial cells from apoptosis and promotes sinusoidal regeneration^39^. Of particular relevance, biochemical studies in C2C12 myoblasts demonstrated that BRM associates with the Notch-responsive *Hes1* promoter in a histone acetylation-associated state even in the absence of Notch activation^45^, suggesting that the *Hes1* promoter may be maintained in a transcriptionally poised state. Our findings raise the possibility that a related mechanism operates in SECs, whereby BRM establishes or maintains transcriptional competence at *Hes1* before injury, allowing the endothelium to rapidly engage a regenerative Notch program upon stress. Although direct BRM occupancy and chromatin remodeling at the *Hes1* locus remain to be established in SECs, this model provides a potential mechanism by which BRM -dependent chromatin regulation could prime niche cells for future stress responses.

The consequences of impaired niche regeneration extend beyond vascular structure to the spatial organization of hematopoiesis. The increased HSC-to-sinusoid distance observed during regeneration in BrmKO mice is consistent with previous work showing that failure to restore endothelial Jag2 with aging is associated with dissociation of HSCs from sinusoids^24^. More recently, an age-associated shift of Jag2 from trans-activation to cis-inhibition was shown to impair HSC polarity, asymmetric division, and regenerative capacity^37^. Our findings extend this model upstream by identifying BRM as a regulator of the niche environment that supports Jag2–Notch signaling and the normal spatial relationship between HSCs and sinusoidal vessels.

The close resemblance between regenerative failure and physiological aging suggests a broader implication of our findings. The Brm-deficient microenvironment recapitulated several features of the aged bone marrow niche, including sinusoidal dilation, impaired Notch signaling, altered HSC positioning, and myeloid-biased hematopoiesis. Moreover, BRM expression declines with physiological aging in SECs. These observations raise the possibility that niche aging arises, at least in part, from a progressive loss of regenerative capacity rather than solely from time-dependent deterioration. We propose that age-associated BRM decline reduces the ability of niche cells to fully recover from repeated physiological insults. Although defects arising from individual episodes may be transient, incomplete restoration over repeated cycles of stress and repair throughout life could progressively accumulate, ultimately converting impaired stress adaptation into persistent structural and functional deterioration of the bone marrow niche.

This model also provides a framework for understanding why the effects of BRM deficiency differ among vascular niche populations. SECs, which undergo extensive remodeling following myelosuppressive injury, showed persistent quantitative and structural abnormalities in the absence of BRM, whereas the effects on AECs were comparatively limited. LepR⁺ stromal cells exhibited a distinct pattern, with relatively preserved homeostasis but an impaired response during acute regeneration. Thus, the requirement for BRM may be greatest in niche populations that undergo dynamic remodeling in response to hematopoietic stress, consistent with a predominant role for BRM in stress adaptation rather than static tissue maintenance.

A limitation of this study is the use of systemic Brm knockout mice, which precludes complete separation of niche-specific effects from hematopoietic cell-intrinsic functions. Nevertheless, transplantation of WT hematopoietic cells into BrmKO recipients demonstrates that BRM deficiency within the recipient microenvironment is sufficient to impair hematopoietic reconstitution and promote a myeloid-biased progenitor composition. Future studies using endothelial- and stromal-specific Brm conditional knockout models will be required to define the relative contributions of individual niche populations and to directly test whether progressive loss of BRM within these compartments is sufficient to drive age-associated niche dysfunction. In conclusion, our findings identify BRM as a key regulator of bone marrow niche regenerative resilience. By maintaining Jag2-producing niche populations while preserving Notch2–Hes1 responsiveness in sinusoidal endothelial cells, BRM supports efficient vascular niche regeneration after hematopoietic stress and restoration of an appropriate environment for HSCs. We propose that age-associated decline of this BRM-dependent stress-adaptive program may permit incomplete repair to accumulate over time, providing a mechanistic link between chromatin remodeling, regenerative failure, and bone marrow niche aging.

## Acknowledgments

The authors thank T. Setsu, Y. Sakihama, and T. Shimizu for technical assistance; Drs. T. Arinami and Y. Iijima for providing BrmKO mice; and all members of the laboratory for helpful discussions. This work was supported by JSPS KAKENHI Grant-in-Aid for Transformative Research Areas (A) (21H05254 to R.N.), Scientific Research (B) (26K02192 to R.N.), Scientific Research (C) (24K11539 to E.N.), Grant-in-Aid for JST SPRING (JPMJSP2148 to H.S.), Grant-in-Aid for JST Moonshot Research and Development Program (JPMJMS2024-7 to R.N.), the COI-NEXT Support Unit for Imaging Science at Kento, and by a grant from the Hyogo Science and Technology Association to R.N.

## Authorship

### Contribution

E.N. designed the research; H.S., H.M., N.Y., T. Shiina and E.N. performed the experiments; H.N., S.K., H.E., R.Y., S.U. provided technical assistance; H.S., H.N. and E.N. analyzed the data; H.S. prepared the figures; H.S. and E.N. wrote the manuscript; A.I., T. Suda, R.N. provided resources and research support; and H.M., N.Y., H.N., S.K., H.E., T. Shiina, R.Y., S.U., A.I., T. Suda, and R.N. reviewed the manuscript.

### Correspondence

Eriko Nitta, MD, PhD, Division of Structural Medicine and Anatomy, Kobe University Graduate School of Medicine, 7-5-1 Kusunoki-cho, Chuo-ku, Kobe 650-0017, Japan.

---

**Supplementary Figure 1.**
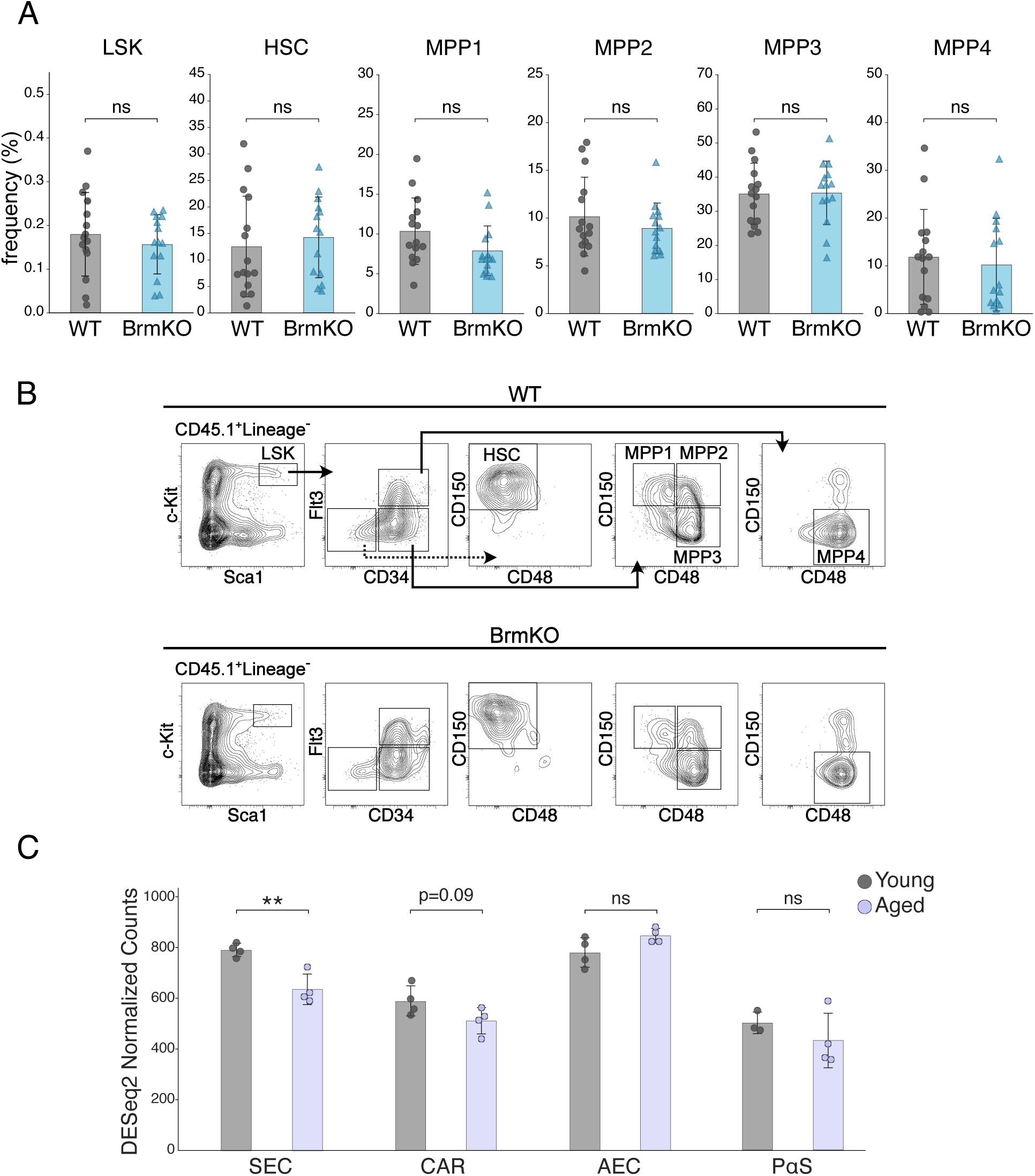
Donor-derived hematopoietic stem and progenitor cells and age-associated BRM decline in the bone marrow niche. (A) Frequencies of donor-derived (CD45.1⁺) LSK cells, HSCs, MPP1, MPP2, MPP3, and MPP4 in the BM of WT and BrmKO recipients at 16 weeks after transplantation (WT, *n* = 15; BrmKO, *n* = 14). Data are pooled from at least 3 independent experiments and presented as mean ± SD. *P* values were calculated using a two-tailed Mann–Whitney U test. ns, not significant. (B) Representative flow cytometry gating plots of donor-derived hematopoietic stem and progenitor cell populations. (C) BRM expression in bone marrow niche populations of adult and aged mice, analyzed from the publicly available RNA-seq dataset GSE133922. Normalized BRM expression is shown for sinusoidal endothelial cells (SECs), CXCL12-abundant reticular (CAR) cells, arteriolar endothelial cells (AECs), and CD140b⁺Sca-1⁺ (PαS) stromal cells isolated from 2-month-old adult (Young, gray) and 2-year-old aged (Aged, purple) male mice. Each dot represents one biological replicate prepared from cells pooled from two mice (*n* = 4 per group; *n* = 3 for Young PαS cells). Bars indicate the mean, and error bars indicate SD. *P* values were calculated using a two-tailed Welch’s *t* test. ns, not significant;\*\**P* = 0.0086.

**Supplementary Figure 2.**
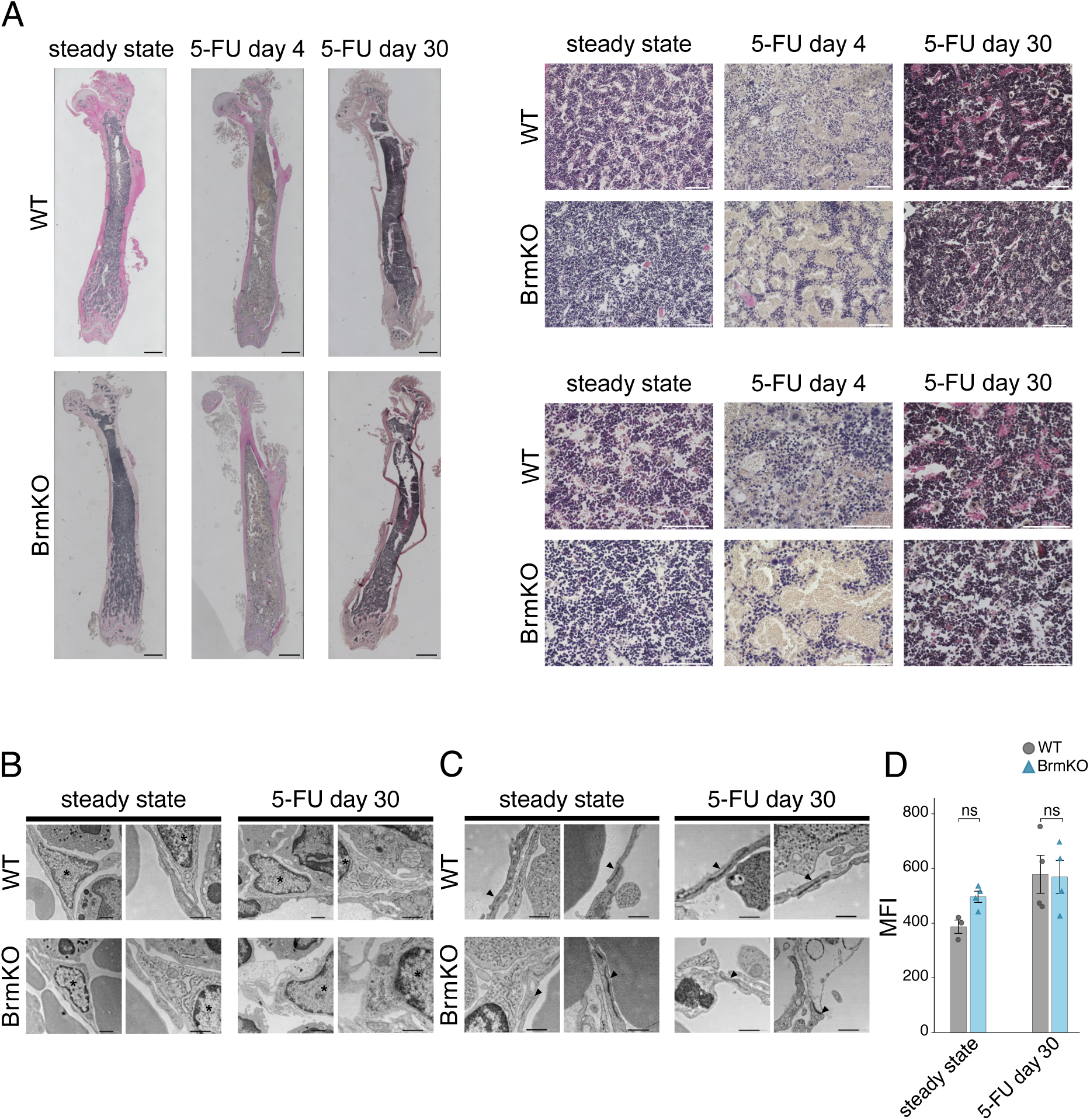
Histological and ultrastructural features of the sinusoidal niche. (A) Representative hematoxylin and eosin (H&E)-stained femur sections from WT and BrmKO mice at steady state and days 4 and 30 after 5-FU treatment. Scale bars, 1000 μm (whole bone view, left) and 100 μm (high-magnification view, right). (B) Representative TEM images of perivascular stromal cells at steady state and day 30 after 5-FU treatment. Scale bars, 1 μm and 500 nm. (C) Representative TEM images of endothelial cell–cell junctions (black arrowheads) at steady state and day 30 after 5-FU treatment. Scale bar, 500 nm. (D) Quantification of mean CD144 (VE-cadherin) fluorescence intensity in CD144⁺ sinusoidal vessel regions of interest from whole-mount immunofluorescence images (*n* = 3–4 mice per group). Data are pooled from at least 3 independent experiments and presented as mean ± SD. *P* values were calculated using a two-tailed Mann–Whitney U test. ns, not significant.

**Supplementary Figure 3.**
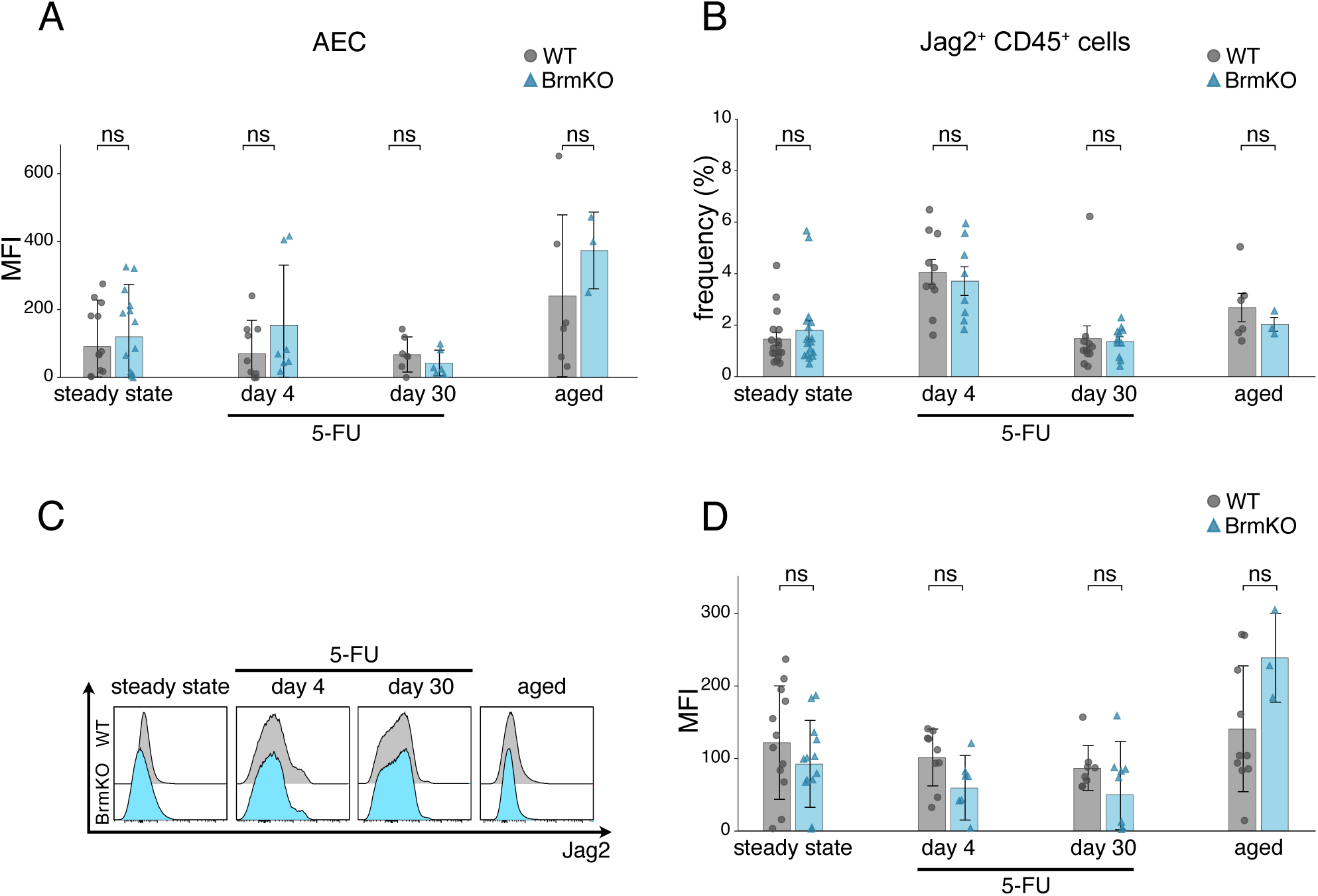
Jag2 expression across bone marrow cell populations. (A) Flow cytometric quantification of Jag2 median fluorescence intensity (MFI) in arteriolar endothelial cells (AECs) at steady state, after 5-FU treatment (days 4 and 30), and in aged mice (*n* = 3–13 per group). (B) Frequency of Jag2⁺CD45⁺ hematopoietic cells in the bone marrow of WT and BrmKO mice at steady state, after 5-FU treatment (days 4 and 30), and in aged mice (*n* = 3–17 per group). (C) Representative flow cytometric histogram overlays of Jag2 expression in CD45⁺ hematopoietic cells. (D) Quantification of Jag2 MFI in CD45⁺ hematopoietic cells (*n* = 3–13 per group). Data are pooled from at least 3 independent experiments and presented as mean ± SD. *P* values were calculated using a two-tailed Mann–Whitney U test. ns, not significant.

